## Supplemental Figures for "Transient intracellular acidification regulates the core transcriptional heat shock response"

*Supporting information for 'Transient intracellular acidification regulates the core transcriptional heat shock response', Triandafillou et al. 2019*

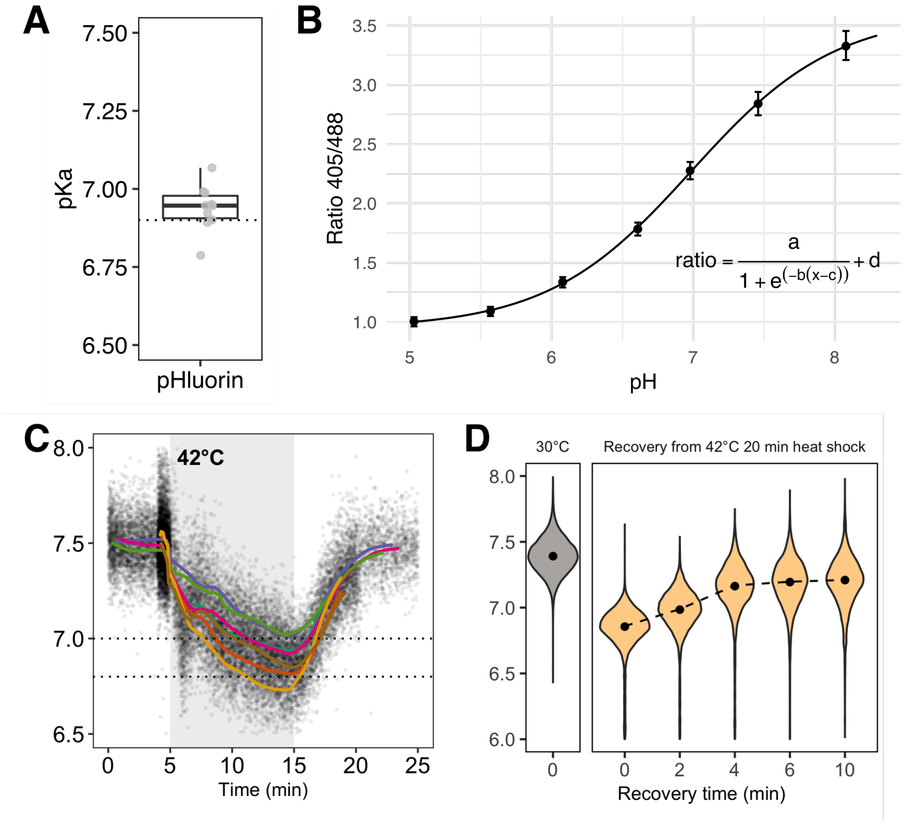


**Figure S1. Measurement of intracellular pH during stress. (A)** Comparison between calibration curves taken on different days. The curves are compared by solving for the apparent $pK_a$ of pHluorin (see Methods section for equation); while absolute values of the ratio vary by day and instrument, the $pK_a$ should be constant. The in vitro $pK_a$ as calculated in Bagar et al. (2009) is shown with the dashed line. Each point is a separate experiment, n=10. See Methods for full details. **(B)** A representative pHluorin calibration curve showing the relationship between intracellular pH and fluorescence ratio. Error bars are the standard deviation of the population of cells measured. **(C)** Traces of intracellular pH as a function of time in cells expressing pHluorin and perturbed with a 42°C, 10 minute heat stress. Each point is an observation of a single cell; colored lines are the moving average of one experiment. Individual points are subsampled for clarity; each experiment has at least 10 000 cells. **(D)** Intracellular pH drops in response to a 42°C, 20 minute heat stress; the degree of acidification is the same as a 10 minute heat stress.

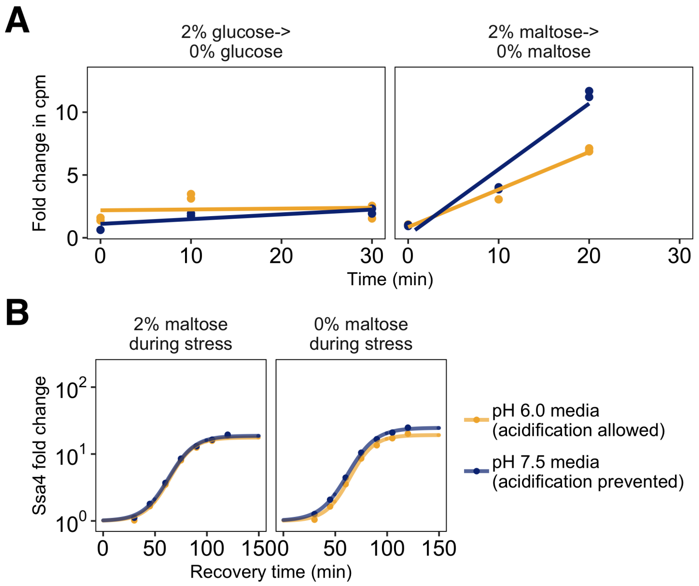


**Figure S2. Preventing stress-associated acidification delays or impairs the heat shock response when translation is inhibited. (A)** Measurement of the incorporation of radiolabeled amino acids into total cellular protein as a function of time after transfer to sugar-free medium. Translation abruptly ceases after withdrawal of glucose (left hand side), but continues after maltose withdrawal (right hand side). In both cases, acidification does not affect the translation rate. **(B)** Induction of Ssa4-mCherry for cells stressed after growth in maltose (left hand side) or growth in maltose followed by brief maltose withdrawal (right hand side). Yellow curves are data from cells in acidic media where acidification is prevented, blue are data from cells grown in media at the resting pH where acidification is prevented. Ssa4 induction after maltose withdrawal is pH-independent, demonstrating that translation collapse rather than nutrient withdrawal explains the difference between induction in the right hand side of **B**.

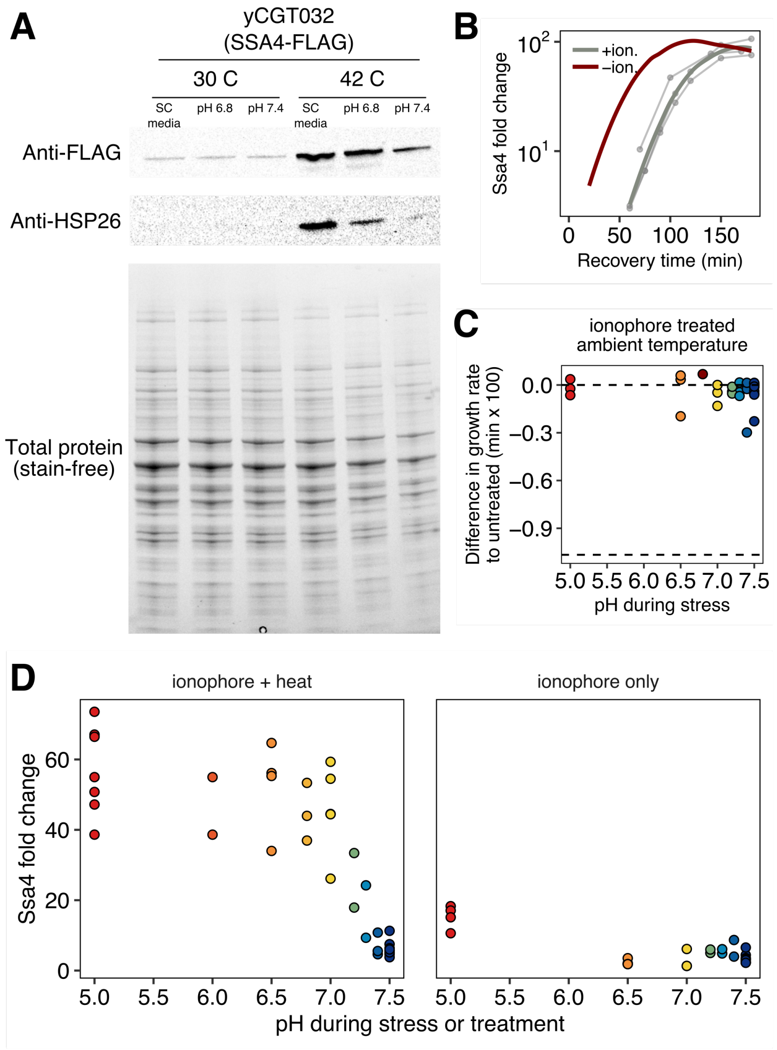


**Figure S3. Quantitative control of intracellular pH using an ionophore. (A)** Western blot and total protein gels for yeast carrying a the genomic copy of SSA4 tagged with a FLAG tag heat stressed with and without ionophore treatment, as described in Growth Conditions in the Methods section. Samples were taken 1 hour after stress. **(B)** Induction of Ssa4 during recovery from normal (red) or pH-manipulated (gray, pH 6.8) stress. Thin curves are individual experiments and thick curves are smoothed conditional means (see Methods for details). The red curve is the same data from Figure 1D for comparison. Although pH manipulation causes a delay in Ssa4 production, it does not affect the ultimate level of induction. **(C)** Growth rate difference in cells treated with ionophore for 35 minutes at room temperature followed by return to ambient growth conditions. Competitor was untreated cells. The values cluster around zero, indicating little to no loss of fitness due to ionophore treatment. Bottom dashed line shows theoretical minimum of the growth rate difference, which would result if cells completely arrested growth. These data are the same as those in Figure 5B, light-colored points. **(D)** Comparison of Ssa4-mCherry induction in ionophore treated cells that were either heat stressed (left) or held at room temperature (right). Acidification artificially induced by ionophore treatment does not cause appreciable accumulation of stress protein.

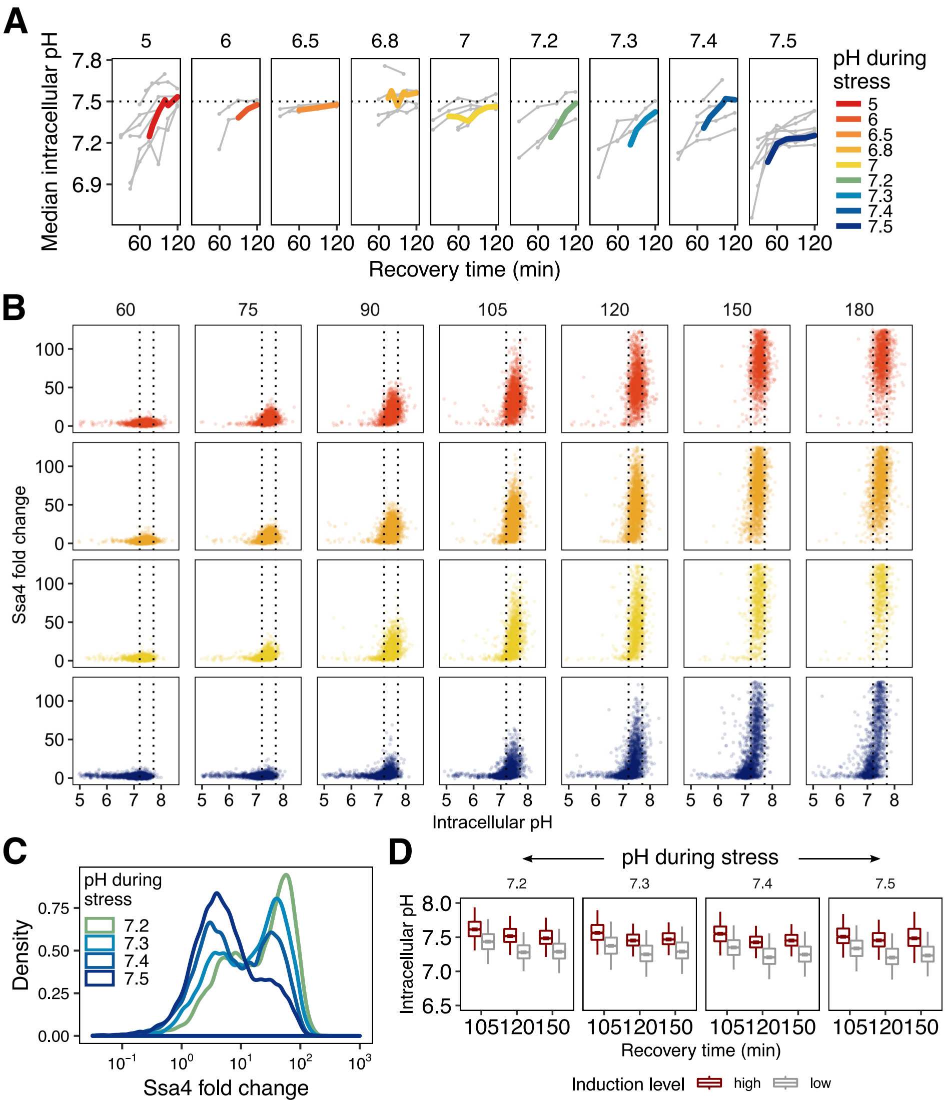


**Figure S4. Post-stress acidification can rescue induction of the heat shock response.** **(A)** Intracellular pH recovery after stress at different intracellular pHs. Thick, colored lines are the moving average of individual experiments (thin gray lines). **(B)** Recovery of intracellular pH is correlated with high Ssa4 levels on the single-cell level. Cells that are stressed at the resting pH have a large proportion of cells that do not recover intracellular pH and do not produce high levels of Ssa4. **(C)** Bimodal distribution of Ssa4 fold-change in cells stressed close to or at the resting pH. **(D)** Intracellular pH distributions for both high-expressing (red) and low-expressing (gray) cells for all conditions shown in C at multiple timepoints during recovery from heat shock.

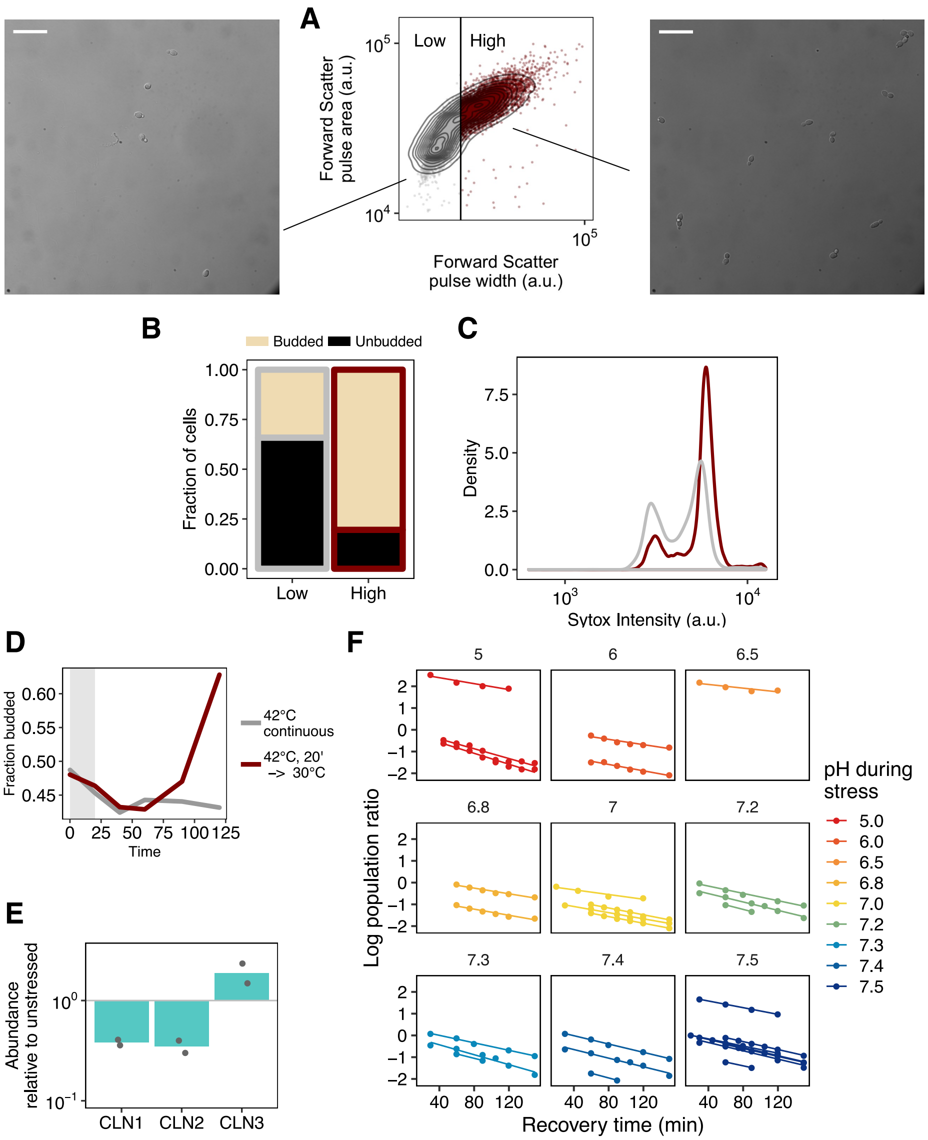


**Figure S5. Fitness, intracellular pH, and heat shock protein production during recovery is correlated in single cells. (A)** Cells were partitioned into two categories in the forward scatter width channel and were sorted based on this partitioning. Sorted cells were fixed and visualized by microscopy. Representative images for each population are shown. Scale bar is 25 $\mu$m. **(B)** Quantification of microscopy data; N = 217 cells scored. **(C)** Fixed cells were stained with Sytox and analyzed by flow cytometry to assess DNA content. The relative heights of the two peaks reflect the proportion of cells in each population that have doubled their DNA, and are thus actively growing. **(D)** Proportion of budded cells as a function of time for cells moved from 30°C and held at 42°C (gray line) or cells that experienced a 42°C heat shock followed by recovery at 30°C (red line). Both populations initially show a dip in the proportion of budded cells, but populations returned to ambient growth temperature then rapidly and synchronously re-enter the cell cycle, as evidenced by an increased in the proportion of budded cells. **(E)** Relative mRNA abundance for three cell cycle transcripts after a 42°C, 20 minute heat stress. Degradation of CLN1 and CLN2 is characteristic of heat shock (Rowley et al., 1993). **(F)** Fits used to determine the relative growth rate for all data shown in Figure 5B. The log of the population ratio as a function of time was fit with a line using linear least squares.

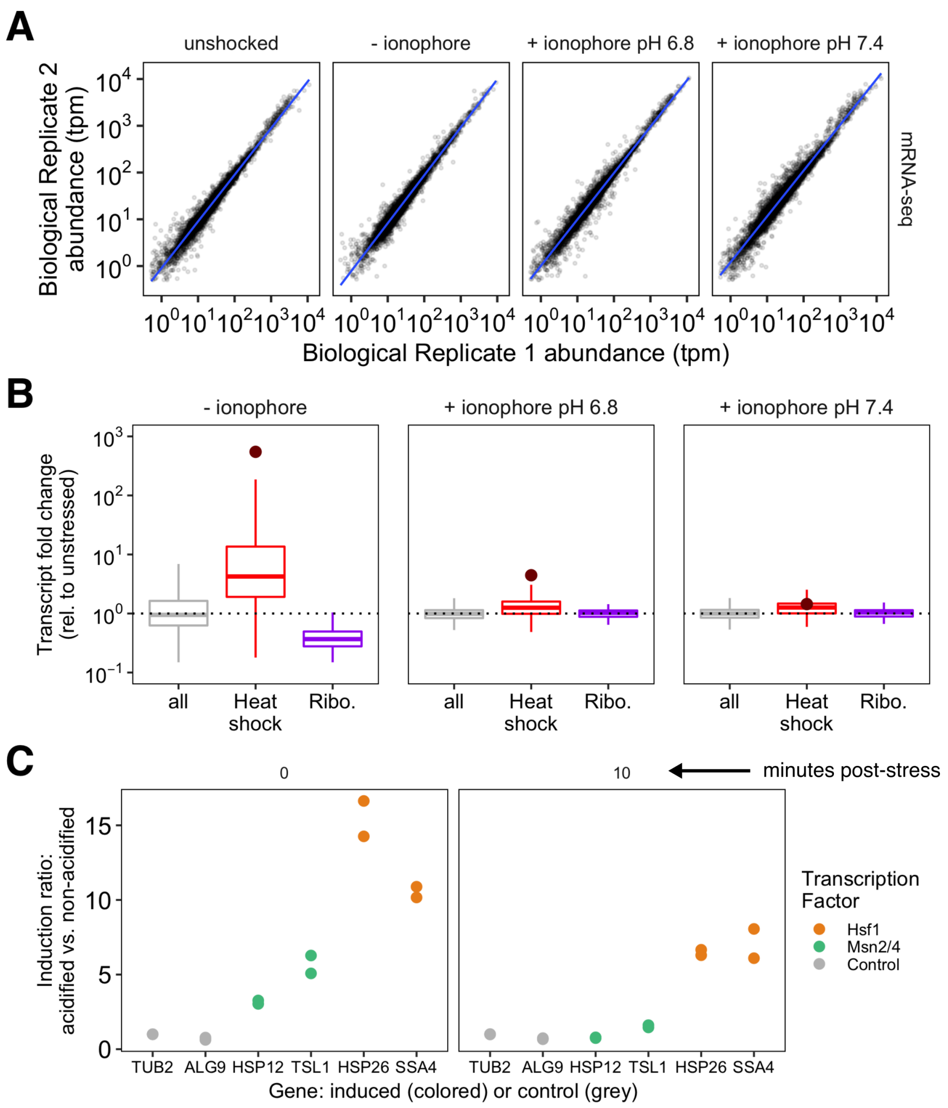


**Figure S6. Failure to acidify during stress specifically represses Hsf1-activated genes.** **(A)** Correlation between mRNA-Seq biological replicates. **(B)** Induction (fold change in stressed cells relative to unstressed cells) for different pHs during stress and different categories of genes. **(C)** qPCR validation of the transcription factor specificity demonstrated in the sequencing experiment in 6C.

### Competing Interests

The authors declare no competing interests.

### Acknowledgements

Research reported in this publication was supported by the National Institute of Biomedical Imaging And Bioengineering of the National Institutes of Health (NIH) under Award Number T32EB009412, and the National Science Foundation Graduate Research Fellowship under Grant No. DGE-1144082. CDK was supported by the NIH award number T32 GM007183. ARD acknowledges support from the NIH, award number R01 GM109455. DAD acknowledges support from the NIH, award numbers R01 GM126547 and R01GM127406, and from the US Army Research Office, award number W911NF-14-1-0411. The authors thank the University of Chicago Flow Cytometry Core for help with flow cytometry data collection and the Functional Genomics Core at the University of Chicago for assistance with sequencing. The authors also acknowledge members of the Drummond and Dinner labs for helpful comments and discussions.
